# The parasite *Schistocephalus solidus* secretes proteins with putative host manipulation functions

**DOI:** 10.1101/2020.02.03.932509

**Authors:** Chloé Suzanne Berger, Jérôme Laroche, Halim Maaroufi, Hélène Martin, Kyung-Mee Moon, Christian R. Landry, Leonard J. Foster, Nadia Aubin-Horth

## Abstract

**Background:** Manipulative parasites are thought to liberate molecules in their external environment acting as manipulation factors with biological functions implicated in their host’s physiological and behavioural alterations. These manipulation factors are part of a complex mixture called the secretome. While the secretomes of various parasites have been described, there is very little data for a putative manipulative parasite. It is necessary to study the molecular interaction between a manipulative parasite and its host to better understand how such alterations evolve.

**Methods:** Here, we used proteomics to characterize the secretome of a model cestode with a complex life cycle based on trophic transmission. We studied *Schistocephalus solidus* during the life stage in which behavioural changes take place in its obligatory intermediate fish host, the threespine stickleback (*Gasterosteus aculeatus*). We produced a novel genome sequence and assembly of *S. solidus* to improve protein coding gene prediction and annotation for this parasite. We then described the whole worm’s proteome and its secretome during fish host infection using LC-MS/MS.

**Results:** A total of 2 290 proteins were detected in the proteome of *S. solidus*, and 30 additional proteins were detected specifically in the secretome. We found that the secretome contains proteases, proteins with neural and immune functions, as well as proteins involved in cell communication. We detected Receptor-type tyrosine-protein phosphatases, which were reported in other parasitic systems to be manipulation factors. We also detected 12 *S. solidus*-specific proteins in the secretome that may play important roles in host-parasite interactions.

**Conclusions:** Our results suggest that *S. solidus* liberates molecules with putative host manipulation functions in the host and that many of them are species specific.

## BACKGROUND

Parasites have major impacts on their hosts, including on their morphology (1), physiology (2) and behaviour (3, 4). To induce these complex changes in their hosts, it has been proposed that parasites produce, store and release manipulation factors that interfere with the host physiological and central nervous systems (5–7). These manipulation factors are thought to be part of a complex mixture of molecules called the secretome, which is a key element of parasite-host interactions (6). The secretome of a parasite includes lipids (8), nucleic acids (9) and proteins (10), which are sometimes protected inside extracellular vesicles (11). Using molecular and bioinformatics approaches, the proteomic fraction of secretomes of parasites infecting humans (12) and livestock (13) have been described, both in terms of protein composition and function (14, 15) (see Table 1 for a review).

**Table 1.**
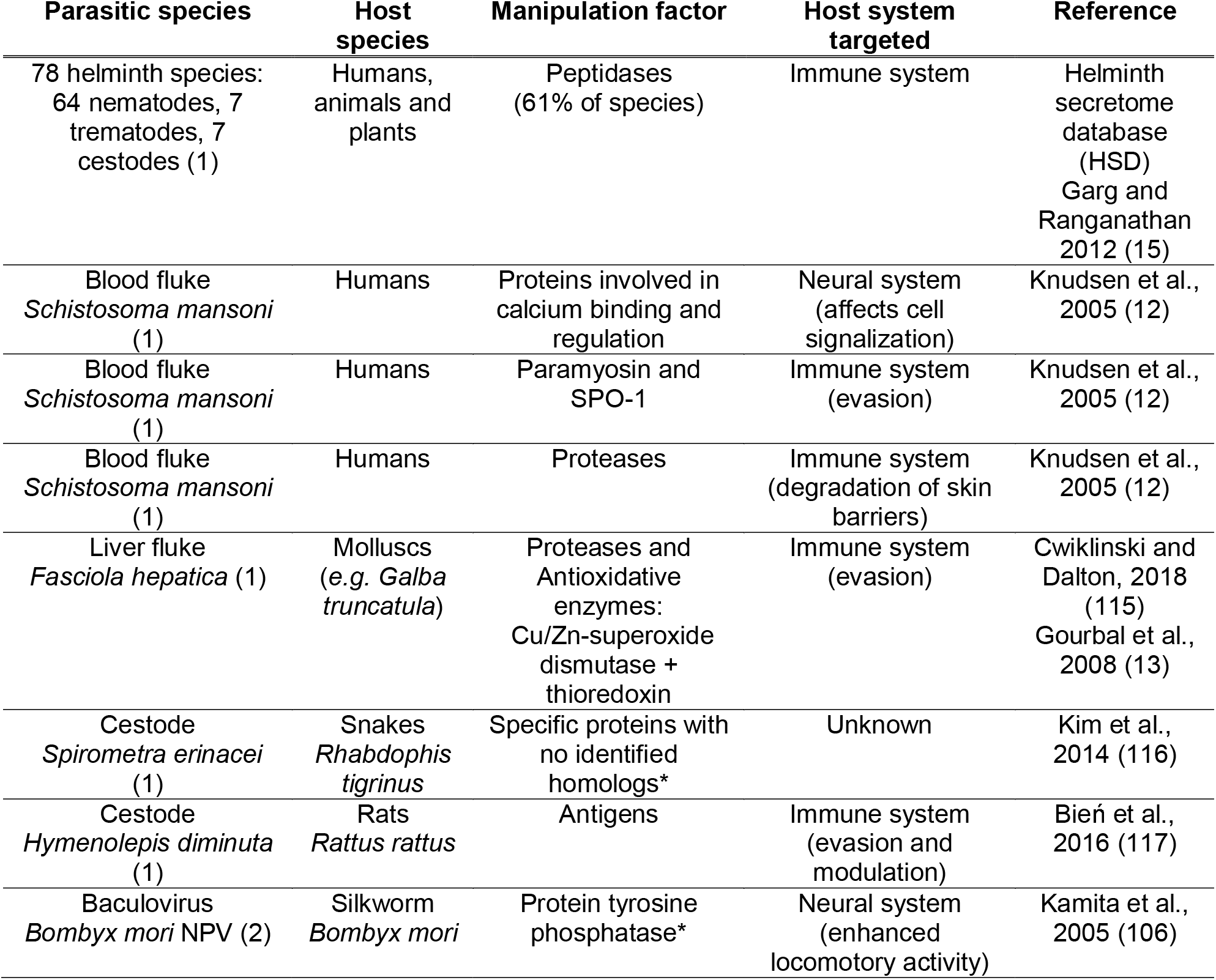
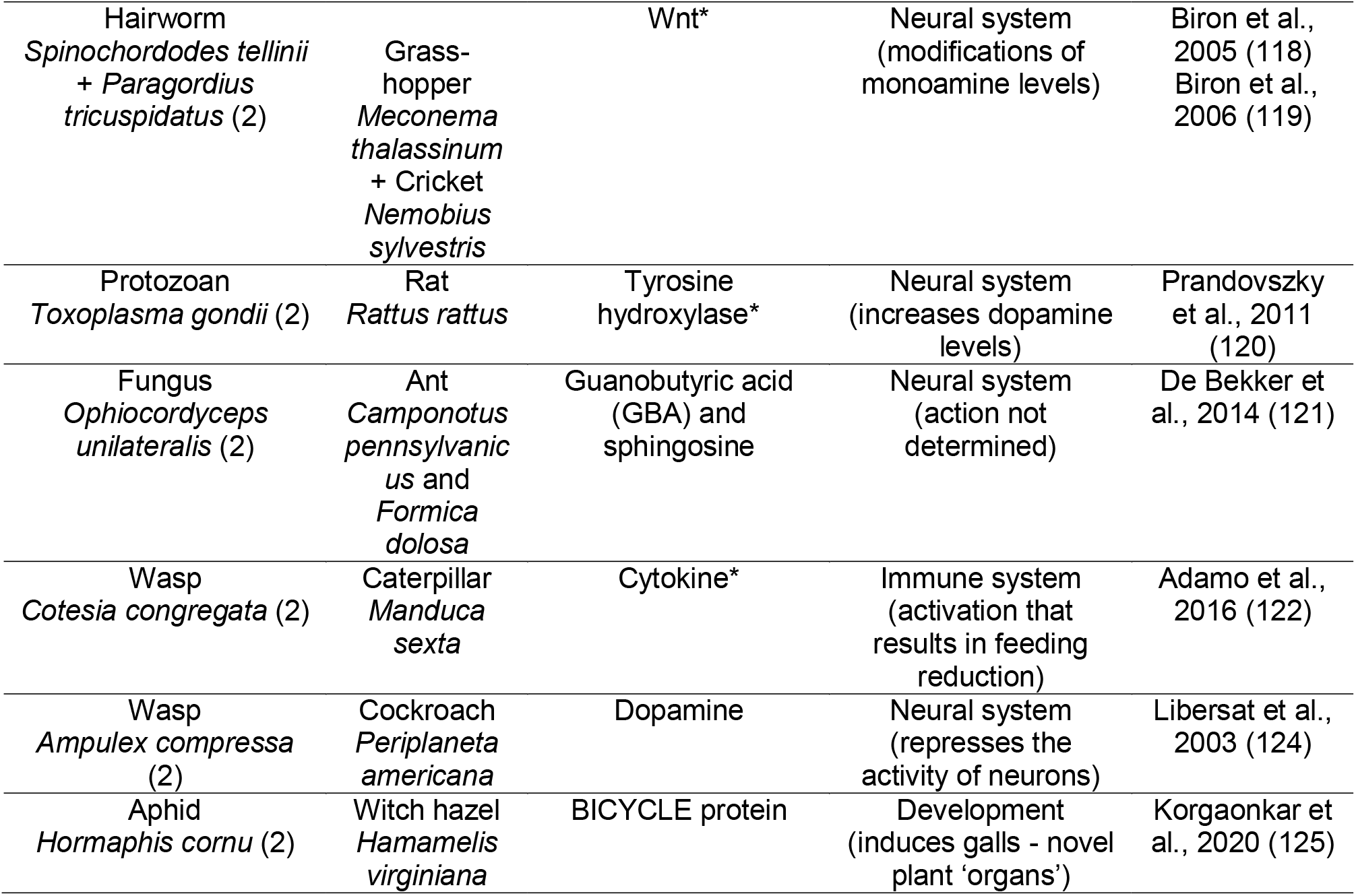
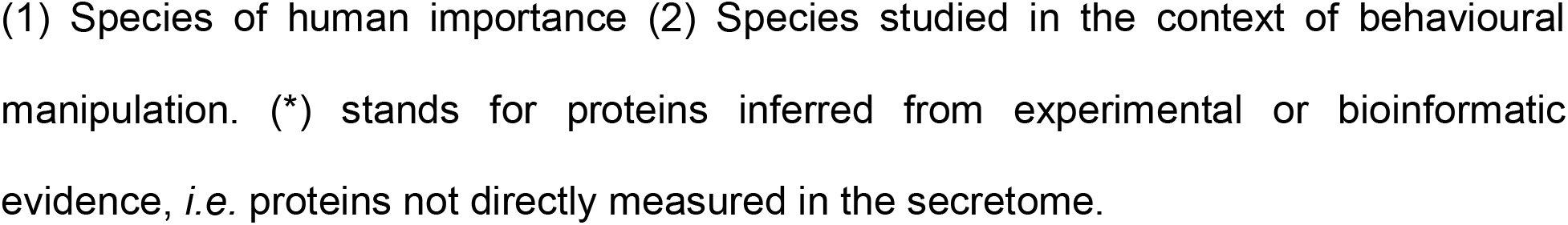
Proteomic content (directly measured or inferred) of the secretomes of different parasite species.

The secretomes that have been examined so far are enriched in peptidases and proteases (12, 15), which are known to weaken the host immunity barriers. Other secreted proteins, such as paramyosin in the blood fluke *Schistosoma mansoni*, have been shown to help the parasite to escape the host immune response, while secreted proteins involved in calcium functions have impacts on the host neural activity (12). In the context of behavioural manipulation, the secretome is a logical potential source of manipulation factors. However, the secretome content of a behaviour-manipulating parasite has rarely been investigated, to the point that secretomes are referred to as “the missing link in parasite manipulation” (7). The literature contains several reports from which it is possible to infer a list of putative manipulation factors, which would target the neural and the immune systems of the hosts and induce behavioural changes (Table 1). Our knowledge regarding if and how many proteins with neural and immune functions can be found in the secretomes of manipulative parasites is very limited, and is based in many cases on inferred proteins rather than actual detection.

One particularly powerful model to study behavioural manipulation is the cestode *Schistocephalus solidus* (16). This parasite exhibits a complex life cycle based on trophic transmission that includes three hosts: a copepod, the threespine stickleback (*Gasterosteus aculeatus*, obligatory intermediate fish host) and a fish-eating bird, in which *S. solidus* reproduces (16, 17). *S. solidus* infects the threespine stickleback’s abdominal cavity through the ingestion of a parasitized copepod (18). The consequences of the infection by *S. solidus* on the threespine stickleback’s morphology (19), physiology (20), immune system (21), and behaviour (22) are well-documented. For example, sticklebacks infected by *S. solidus* show drastic behavioural changes that result in a loss of the anti-predator response (16): infected fish are more exploratory (23), less anxious (24) and bolder in the presence of a predator (25) than non-infected fish.

Most of these behavioural alterations seen in infected fish appear after several weeks, when the worm has grown to reach the infective stage within its intermediate host (*i.e*. larger than 50 mg) (26). The infective stage coincides with the time at which *S. solidus* is ready to reproduce in the bird (16, 27), which also generally corresponds to the activation of the immune response in the host. In the first phase of infection, the adaptive immune response is generally not activated in the fish. It is only when the worm reaches the infective stage that an ineffective up-regulation of the respiratory burst activity occurs (21). Nevertheless, activation of the immune response through the production of reactive oxygen species (ROS) by granulocytes has been shown to occur early during infection, depending on the genotype of the stickleback population (126). Several studies have suggested that the manipulation of the stickleback’s behaviour increases the worm’s transmission rate to its final avian host (16). Yet, the adaptive value for *S. solidus* of such behavioural changes has never been demonstrated (28), and it is possible that these behavioural modifications in the fish may solely result from a side effect of infection (7), such as the effect of the parasite mass burden or of the activation of the host immune response (24, 109). To demonstrate that behavioural changes in the host are the result of direct parasitic manipulation, the first step requires to determine if the parasite can liberate molecules in its external environment, and if yes, to study their functions in relation with the host’s phenotype perturbations.

This host-parasite system was used for the first experimental demonstration of a complex life cycle of a trophically transmitted parasite in 1790 (as reviewed in (16)). A rich body of work on this model system during the past 50 years has shown, using enzymatic assays, that the activity of proteases (29) and transferases (30) are required for *S. solidus* survival and growth. Furthermore, a partial genome of the worm is available (31) and extensive transcriptome data has been produced (32, 33). Quantification of the transcriptome dynamics across the life stages has uncovered that when the worm reaches the infective stage in its fish host, genes involved in neural pathways and sensory perception have higher expression levels, compared to the earlier stages in the same host, which are characterized by upregulation of growth-related pathways (33). Furthermore, vesicles are present inside the *S. solidus* tegument, as shown through scanning and transmission electron microscopy (34). Moreover, *S. solidus* excretes (through passive mechanisms) or secretes (through active mechanisms) molecules, including (uncharacterized) proteins, and these secretions are sufficient to affect its fish host behaviour (35). Finally, a well-annotated genome of the threespine stickleback host is also available (36), which is important to adequately differentiate proteins coming from the host and the parasite. The *Schistocephalus*-stickleback system is thus appropriate to test the presence of manipulation factors. However, the nature of the protein content of the *S. solidus* secretome has never been explored (7, 35). Analyzing the proteomic fraction of the secretome of *S. solidus,* and its potential enrichment in manipulation factors involved in neuronal and immune functions, will help us to understand if the behavioural changes of the host could be induced by parasitic manipulation through the secretome.

Here, we characterized the worm’s whole-body proteome and the protein fraction of the secretome of *S. solidus* using mass spectrometry (LC-MS/MS) (37). We focused on individuals in the infective stage of development so that the secretome may include manipulation factors that could be associated with the fish host’s behavioural changes. In helminth parasites, a portion of proteins from the proteome are passively released in the external environment as metabolic waste products, contributing to an important fraction of their secretome content (38). Therefore, the secretome is generally a subset of the proteome in terms of protein content (39). We thus expected that the secretome of *S. solidus* would also mainly be a subset of its proteome. However, in the context of behavioural manipulation, we hypothesized that proteins could also be actively liberated by the parasite in its external environment (38), so that they would be enriched in the secretome compared to the proteome. Based on what has been described in previous parasitic systems (references reviewed in Table 1), if *S. solidus* manipulates stickleback behaviour with its secretions as we hypothesize, then its secretome would include proteases, as well as proteins with neural and immune functions. Because the worm is not in direct contact with the vertebrate host brain (16), we also expected to detect proteins involved in cell communication, cell-cell signaling or transport functions. These proteins would mediate the communication of the worm with its host brain to induce potential neural and immune changes, and ultimately behavioural alterations.

As the genome of *S. solidus* is used during LC-MS/MS as a reference database, a more thorough annotation than the one publicly available (31) of the genome could allow us to detect more proteins, including potential manipulation factors. We therefore used a multi-pronged approach combining genomics with proteomics (described in Figure 1). We first sequenced the genome of *S. solidus* using a combination of long and short reads to combine longer contigs and improved annotation. Then, we investigated the global proteomic composition of the proteome and secretome of *S. solidus* using LC-MS/MS.

**Figure 1.**
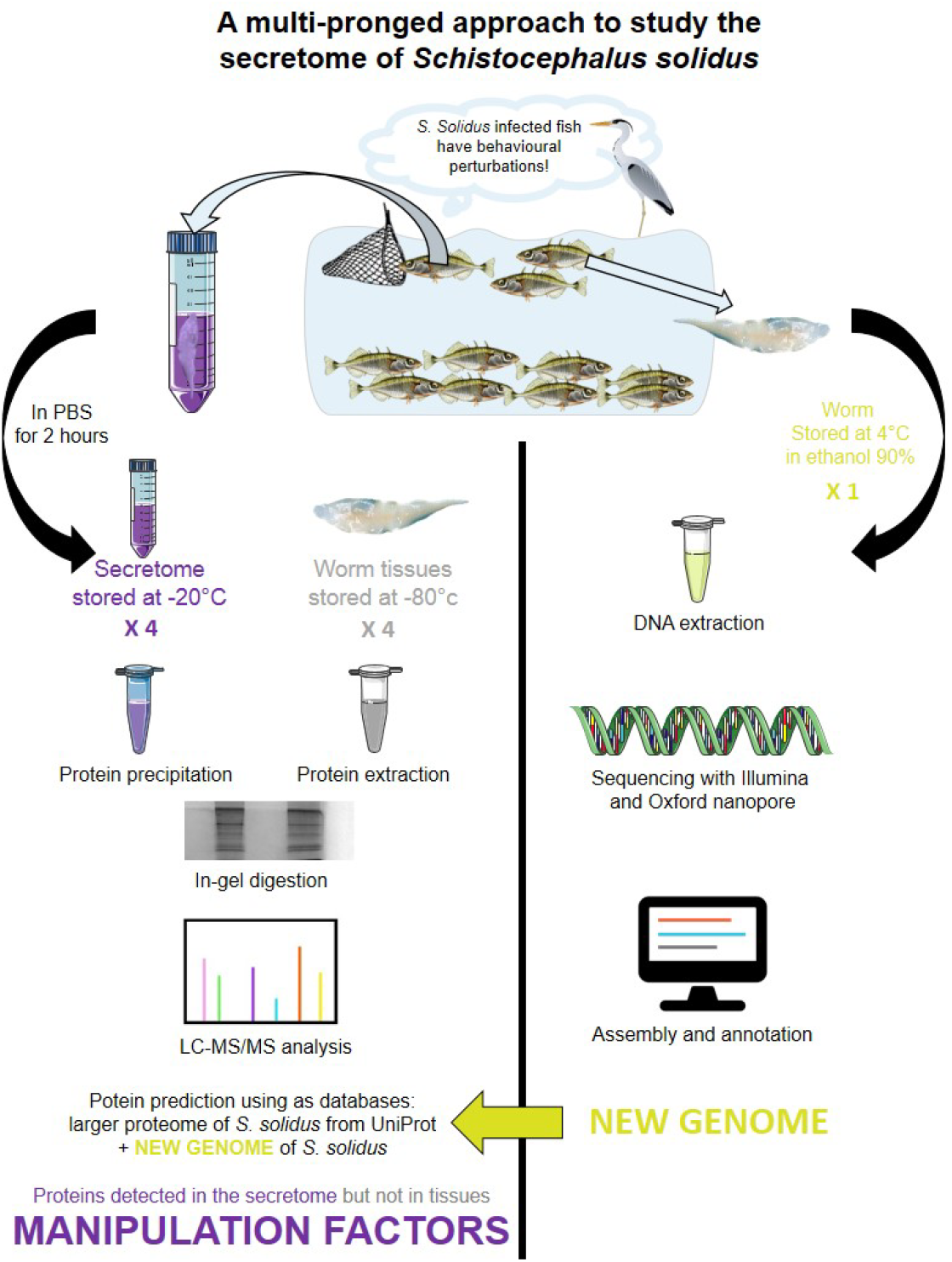
The multi-pronged approach designed to detect potential manipulation factors in the secretome of *Schistocephalus solidus*. On the left, the proteomic approach based on LC-MS/MS aims at describing the global proteomic composition of the proteome and the secretome of *S. solidus*. On the right, the genomic approach aims at producing a novel genome sequence and assembly of *S. solidus* to improve protein coding gene prediction and annotation for this parasite. The genome, whose quality is improved, can be used as a new reference database to infer proteins in the proteome and the secretome of *S. solidus*. Combined together, these approaches allow to characterize the secretome of *S. solidus* at the infective stage, including the uncovering of proteins detected only in that fraction and not in its proteome, thus representing potential manipulation factors.

## METHODS

### Genome sequencing of Schistocephalus solidus

#### Worm collection

We caught threespine sticklebacks from Lac Témiscouata (Québec, 47°40′33″N 68°50′ O) where fish are known to be infected by *S. solidus* (24), using minnow traps in July 2015. Fish were brought to the “Laboratoire Aquatique de Recherche en Sciences Environnementales et Médicales” at Université Laval (Québec, Canada) and were maintained under a 12h:12h Light:Dark cycle and a water temperature of 15°C. Fish were fed daily with a mixture of blood worms and Artemia. After 10 months, one fish (out of 40) exhibited the morphological changes typically induced by *S. solidus* (19) and was consequently sacrificed to collect the worm. The fish was euthanized with an overdose of MS-222 (75 mg/L mg/kg) and dissected to confirm the infection by *S. solidus*. The worm was immediately put in ethanol 90% and stored at 4°C until genome sequencing. All the other fish were later used during a behavioural experiment (24).

#### DNA extraction and sequencing

Genomic DNA was extracted using DNeasy Blood and Tissue kit (Qiagen Inc., Valencia, CA, USA) from ∼20 mg of tissues. After lysis, we added 4 µL of RNase A (10 mg/mL). Elution was done twice in 100 µL of elution buffer. To reach the desired concentration for the Nanopore library preparation and Illumina sequencing, we concentrated DNA with a SpeedVac Concentrator (ThermoFisher, Waltham, MA, USA) for one hour.

#### Illumina HiseqX sequencing

The DNA libraries for Illumina sequencing were prepared using a Shotgun PCR-free library preparation (Lucigen) Illumina Library at the McGill University and Genome Quebec Innovation Center (Montréal, Canada). Sequencing was performed at the same center on an Illumina HiSeqX instrument, using paired-ends reads (PE150). A total of 157 475 128 reads were obtained.The estimated genome size is 540 MB (31).

#### Oxford Nanopore Technonologies (MinION) sequencing

Library preparation for *Oxford Nanopore* sequencing was done with a PCR-free ligation sequencing kit SQK-LSK108 (ONT, Oxford, UK). Briefly, approximately 2.5 µg of high molecular weight DNA was repaired using the NEBNext FFPE Repair Mix (NEB, Ipswich, Ma, USA) for 15 min at 20°C before purification with Ampure XP beads. The repaired DNA was end-prepped with the NEBNext Ultra II End Repair/dA-Tailling Module (NEB, Ipswich, Ma, USA), for 30 min at 20°C and 30 min at 65 °C, according to the manufacturer’s instructions and purified with Ampure XP beads. Adapter mix (ONT, Oxford, UK) and Blunt/TA Ligation Master Mix (NEB, Ipswich, Ma, USA) were added to the purified end-prepped DNA and incubated for 15 min at room temperature. Library was then purified with Ampure XP beads and ABB wash buffer and recovered in Elution Buffer (ONT, Oxford, UK). Approximately 333 ng of the purified library was loaded onto a primed R9.4 SpotOn Flow cell (FLO-MIN106), along with RBF (ONT, Oxford, UK) and Library Loading Beads (ONT, Oxford, UK). Sequencing was performed with a MinION Mk1B sequencer running for 48 hours and the MinKNOW software (provided by ONT, Oxford, UK) was used to control the sequencing process. Four libraries were sequenced using this protocol. Base calling was performed with albacore (read_fast5_basecaller.py, ONT Albacore Sequencing Pipeline Software v2.3.1) on all fast5 files. A total of 4 636 932 read sequences resulted from the base calling step for a total of 14 535 522 365 nucleotides.

#### Genome assembly and annotation

All fastq ONT files were pooled in one file prior to assembly. ONT sequences were assembled using Flye v2.4 (40, 41). The final assembly produced 17 882 scaffolds, the largest scaffold being 919 337 nt, with a N50 of 121 189 nt. The mean coverage across scaffolds was 20X. A first phase of correction (polishing) was carried out with nanopolish (v0.10.2) (https://github.com/jts/nanopolish). A total of 3 473 881 changes were applied. A second correction phase with one Illumina HiSeqX paired-end sequence library (20X coverage) was also done with Pilon (v1.22) (42). A total of 4 889 938 changes helped to improve the quality of the assembly sequences. Scaffolds that had an average coverage of less than 10X coverage and those that were shorter than 500 bp were removed because they were of limited use prior to genome annotation. These represented less than 1% of the data and all contigs that contained hits with the transcripts from S. solidus (33) remained after this selection. This left a total of 15 357 scaffolds and 625 207 408 nucleotides. This Whole Genome Shotgun project has been deposited at DDBJ/ENA/GenBank under the accession WEDG00000000.1 and the assembly ID ASM1759139v1.

#### Completeness of the genome assembly

We used a dataset of 24 765 transcripts from *S. solidus* that we previously published (33) and mapped them on the *de novo* assembly using GMAP (v2019-03-15) (71) as implemented in the pipeline GAWN v0.3.2 (https://github.com/enormandeau/gawn). An assembly quality analysis was performed using the Benchmarking Universal Single-Copy Orthologs (BUSCO) and BUSCO groups were searched in the metazoa database (43).

#### Protein-coding gene prediction

To find out the proportion of repeated regions in the genome and to get a masked assembly prior to running BRAKER2 to predict protein coding genes, we built a RepeatModeler (1.0.8) database (44) based on the new genome sequence and ran RepeatMasker (4.0.6) based on that database (45).

We used BRAKER2 (46–51) for protein-coding gene prediction using two approaches. The first approach was *ab initio* as it was not based on external data to find Open Reading Frames (ORFs) but only on genes predicted by GeneMark-ES that are selected for training Augustus (46, 47). The second approach used the alignment (bam files) of the transcripts from *S. solidus* (33) on the genome. The two sets of ORFs obtained with the two approaches were merged and duplicate sequences were removed, which allowed to obtain a final number of predicted ORFs. We used a BLAST analysis to quantify the percentage of the *S. solidus* transcriptome we previously published (33) that was found in the genes predicted using Augustus and vice-versa.

#### Identification of sequences specific to the new assembly

The predicted ORFs were locally aligned using BLAST+ (52, 53) against a database of 43 058 protein sequences from *Schistocephalus solidus* obtained from release 230 of NCBI (March 2019, https://www.ncbi.nlm.nih.gov). From the predicted ORFs, we selected those that had no blast match or that had no significant match based on the fact that the length of the alignment was less that 80% of the length of the query or that had less than 90% of identical nucleotides over the length of the query (54). These unmatched sequences that were specific to the assembly were used as one of the reference databases during LC-MS/MS analysis (see below “Protein identification”).

#### Genome annotation

A functional annotation of the predicted ORFs that were obtained with BRAKER2 (46–51) was performed using Hmmer (version 3.3) (55) against PFAM domain database (release 32) (56) and using orthology assignment with eggNOG-mapper (Evolutionary Genealogy of Genes: Non-supervised Orthologous Groups) (version 2.0.1) (57).

### Mass spectrometry characterization of the worm proteome and secretome

#### Worm and secretome collection for mass spectrometry analysis

##### Sampling of experimental individuals

Whole worms were collected from wild-caught fish acclimatized to laboratory conditions. Juvenile sticklebacks came from Lac Témiscouata (Québec, 47°40’33”N 68°50’15”O), the same lake that was used to collect a worm for genome sequencing. Fish were caught using a seine in August 2016. They were brought to the “Laboratoire Aquatique de Recherche en Sciences Environnementales et Médicales” at Université Laval where they were raised for one year in 80 L tanks under a Light:Dark photoperiod of 12 h:12 h and a temperature of 15°C reflecting their natural environment conditions (Québec, Canada). Fish were fed daily with brine shrimps.

##### Collection of proteome and secretome samples

In summer of 2017, 51 fish were individually isolated in 2 L tanks. The day following isolation, fish were injected with 100 µL of PBS (Phosphate Buffered Saline, pH 7.4 Life Technologies) in their abdominal cavity in order to sample their fluids to detect infection by *S. solidus* following the method described in (58). This protocol was repeated the next day. If fish were detected as infected, they were euthanized with an overdose of MS-222 (75 mg/L mg/kg) and dissected to confirm the infection by *S. solidus* and to collect the worm and its secretome. Fish sex, size and mass, and *S. solidus* mass and number in each fish were noted. We found that 5 fish were infected, each harbouring a worm whose weight was above 50 mg (worm 1 = 485.3 mg; worm 2 = 504.1 mg; worm 3 = 286.5 mg; worm 4 = 544.5 mg; worm 5 = 220.9 mg). All the worms used to collect proteome and secretome samples were far above the mass threshold of 50 mg, which is commonly used to define the parasitic infective stage that coincides with the appearance of the behavioural changes in the fish host (16, 26, 27). Furthermore, behavioural changes in fish infected by worms of similar masses were previously reported in the Temiscouata population (35). Therefore, we considered these worms and secretome samples to be representative of the manipulative stage of *S. solidus*. We only selected fish that were infected by a single worm to prevent potential effects of multiple infections on the proteomic content of the secretome.

The worm secretome was collected according to a protocol adapted from (59). Each worm was rinsed with 1 mL of PBS (Phosphate Buffered Saline, pH 7.4 Life Technologies) to remove fish fluids and then immediately put in a 2 mL tube of PBS. The tube was covered with aluminium foil to protect the worm from light and placed in a water recipient at the same temperature as the fish tanks (15°C) for 2 hours. The worm was removed from the tube and a tablet of Complete, Mini Protease Inhibitor (Sigma) was added to the tube to protect the proteins from the protease activity. The worm tissues were then snap frozen in liquid nitrogen and stored dried at −80°C for proteomics analysis. The liquid in which the worm was incubated (PBS + potential secretome collected) was stored at −20°C. Further experiments were therefore conducted with 5 worms and their respective secretome.

#### Preparation of worm tissues and secretome for in-gel digestion

Worms were individually washed with 2 mL of PBS (Phosphate Buffered Saline, pH 7.4 Life Technologies) to remove potential remaining contaminants from the fish. They were then cut into three equal pieces that were individually put in a tube of 700 µL of lysis buffer (4% (w/v) SDS in 100 mM Tris pH 8 - 10 mM Dithiothreitol DTT). Six sterile ceramic beads (size 2.8 mm) were added to each lysis tube and the samples were homogenized for 20 seconds at 6 000 rpm using a Precellys Homogenizer (Bertin Technologies). Homogenization was repeated three times. Samples were put on ice for 1 min between each run. Samples were then spun at 10 000 x g at 4°C for 10 min. For each individual worm, the three homogenates obtained were pooled together and redistributed into equal volumes into 2 tubes. Samples were heated at 95°C for 10 min, then spun for 10 min at room temperature. The supernatant of each sample was collected into a new tube and protein concentration was calculated using the extinction coefficient measured with a Nanodrop 1000 spectrophotometer (A280 nm; Thermo Scientific). The concentration obtained for each worm lysate was respectively: 24.14 mg/mL; 22.03 mg/mL; 15.66 mg/mL; 28.53 mg/mL; and 11.37 mg/mL. Worm lysates were kept at −20°C before performing in-gel digestion.

For the secretome, the protein concentration of each of the liquids in which the worms were incubated (“secretome” samples) was measured using a Pierce Coomassie (Bradford) Protein Assay Kit (Fisher Scientific). Bovine Serum Albumin (BSA) at 2 mg/mL was used as standard (7-point standard curve ranging from 0 to 25 µg/mL). The concentrations obtained were: 13.11 µg/mL; 15.09 µg/mL; 9.83 µg/mL; 16.59 µg/mL; and 7.46 µg/mL. For each secretome sample, 10 µg of proteins were precipitated with Trichloroacetic Acid (TCA) (60). Precipitated samples were directly used for in gel-digestion.

#### In-gel digestion

For each sample, 50 µg of proteins from worm lysate and 10 µg of proteins from secretome were resolved on a 12% SDS-PAGE with a BenchMark protein ladder (Invitrogen) and a negative control (SDS-PAGE loading buffer and water). Migration was performed during 60 min at 175 V for the worm lysates, and during 30 min at 175 V for secretomes. Coomassie blue G250 (PMID: 15174055) was used for staining overnight (Additional file 1). Each migration lane was cut into 5 fractions for worm lysates, and 3 fractions for secretomes. In-gel digestion was performed on these fractions according to a previously developed protocol (61). Briefly, after cutting the gel into slices, proteins were reduced using 10 mM Dithiothreitol (DTT) for 45 min at 56°C, then alkylated with 55 mM iodoacetamide (IAA) for 30 min at room temperature in the dark. Digestion was performed overnight at 37°C with trypsin (Promega V5113; 0.1-1 µg of trypsin depending on the gel staining intensity). The next day, peptides were extracted using an organic solvent (100 µL acetonitrile) and a step-wise protocol (40% acetonitrile then 100% acetonitrile), and dried (< 50 μL). Following in-gel digestion, a STAGE-TIP protocol using C18 extraction disks (3M Empore) was performed to desalt the samples (62, 63). Samples were acidified with trifluoroacetic acid (TFA) (pH < 2.5) before being passed through the tip. At the end of the STAGE-TIP protocol, samples were completely dried using a Vacufuge Plus concentrator (Eppendorf) and stored at - 20°C until LC-MS/MS analysis.

#### LC-MS/MS analysis

The peptides were run on analyse using a quadrupole time of flight mass spectrometer (Impact II; Bruker Daltonics) coupled to an Easy nano LC 1000 HPLC (Thermo Fisher). More information about the LC-MS/MS analysis can be found in additional information (Additional file 2).

#### Protein identification

We searched the detected mass spectra against the *S. solidus* genome using MaxQuant (version 1.6.1.0) (65). Two searches were independently performed: the first search used the larger proteome of *S. solidus* (43 058 entries, downloaded June 21, 2018 and updated May 6, 2019) from the Universal Protein Resource release 2018-05 and 2019-03 (UniProt https://www.uniprot.org/) as a reference database, which includes proteins predicted from the partial genome of *S. solidus* that was then currently available (31), as well as proteins predicted from the *de novo* transcriptome (32, 33). For the second search, we used the unmatched sequences that we reported to be specific to our genome assembly (36 140 sequences, see Results) as a reference database. The search included common contaminants and variable modifications of methionine oxidation, and N-acetylation of the proteins. The data was filtered for matches passing 1% false discovery rate set by MaxQuant (65). We included the larger proteome of the threespine stickleback host (29 032 entries, downloaded June 21, 2018) from the Universal Protein Resource release 2018-05 (UniProt https://www.uniprot.org/) in the search in order to include proteins originating from the fish host during the MaxQuant data search. Subsequent analyses were performed with all the proteins inferred using the larger proteome of *S. solidus* and the unmatched sequences of the new genome as reference databases.

#### Data analysis of the proteome

Data analysis was performed with Python custom scripts (version 3.6.4) using Jupyter notebooks (version 5.4.0). A template of the custom script is available in additional information (additional file 3). Proteins detected in each worm sample were retrieved using MaxQuant. We filtered out of the dataset protein IDs that were solely attributed to the threespine stickleback (Additional file 4), protein IDs with REV coding (reverse hits for False Discovery Rate filtering) and protein IDs with CON coding (contaminant hits that were added into the search). In some cases, several protein IDs were found by MaxQuant for a specific protein because of high sequence similarities between them.

Research on NCBI (https://www.ncbi.nlm.nih.gov/) and Uniprot (https://www.uniprot.org/) databases demonstrated that the proteins IDs detected for one protein were probably isoforms with identical functions. These multiple protein IDs were kept during annotation to obtain exhaustive functional information, but only the first ID of each protein was kept for enrichment analyses to avoid an over-representation of specific processes or functions (see below). This final dataset was used to describe the global composition of the proteome of *S. solidus*.

We performed two distinct enrichment analyses for the proteins detected in at least one worm sample, and for the proteins detected in all worm samples. For proteins with several protein IDs, we kept only the first ID in order to prevent enrichment bias during analysis. Enrichment analysis was performed using the tool Funrich (version 3.1.3) (66). We constructed a custom reference database using the protein IDs and the Gene Ontology (GO) annotation (biological process, cellular component, molecular function) of all the proteins described in the larger proteome of *S. solidus* available on Uniprot (https://www.uniprot.org/ 43 058 entries). P-values for enrichment were obtained with a hypergeometric test corrected with the Bonferroni method.

#### Data analysis of the secretome

Data analysis was performed with Python using the same approach as for the proteome. We separated proteins into two categories: proteins that were shared between the proteome and the secretome samples, and those that were found only in a secretome sample. For the proteins that were shared between the proteome and the secretome samples, we performed two distinct enrichment analyses: one for the proteins detected in at least one secretome sample and one for the proteins detected in all secretome samples. The enrichment analysis was performed using the tool Funrich (version 3.1.3) (66) using the same approach as for the proteome.

We investigated the annotation of proteins found only in the secretome and not the proteome using a three-step approach. During the first step, the protein ID obtained with the MaxQuant analysis was searched in Uniprot to retrieve its information if available (protein name and sequence, corresponding coding gene, protein function, localization and/or structural information). During the second step, we inferred the function of the protein based on sequence homologies using the Basic Local Alignment Search Tool (BLAST) on NCBI (https://blast.ncbi.nlm.nih.gov/Blast.cgi). We used Blastp (protein-protein) using the Non-redundant protein sequences (nr) database (August 2018) and Tblastn (protein-translated nucleotide) using the Nucleotide collection (nr/nt) database (August 2018). During the last step, when no information was found with the previous steps or to confirm the information previously found, we used the Pfam database (version 32.0) (56) to infer the function of the protein based on domain organization homologies. In some cases, several protein IDs were found by MaxQuant for a specific protein because of high sequence similarities between these IDs. We found by applying the previous approach that in all of these cases, the putatively redundant protein IDs detected for one protein were indeed isoforms with identical functions. These multiple protein IDs were nevertheless kept during annotation (see below) to obtain exhaustive functional information.

### Annotation of proteins

In order to obtain an exhaustive annotation of proteins, we conducted complementary approaches based on sequence and structure analyses (as described below) (67–78) for the proteins detected in the proteome and/or enriched in the secretome for which few or no annotation was available. These analyses were also performed for the proteins that were detected using the unmatched sequences specific to our genome assembly as a reference database, for which limited or no annotation was available.

#### Protein sequence analysis

We predicted secreted proteins using the SignalP-5.0 (classical secretory proteins with option “eukaryota”, http://www.cbs.dtu.dk/services/SignalP/) and SecretomeP 1.0f (non-classical pathway secreted proteins with option “mammalian”, http://www.cbs.dtu.dk/services/SignalP-1.1/#submission) servers (67, 68). In addition, we predicted transmembrane helices in proteins using TMHMM Server v. 2.0 (http://www.cbs.dtu.dk/services/TMHMM-2.0/) (69). The determination of protein domains was performed using HmmScan (http://hmmer.org/) against profile-HMM databases such as Pfam, Gene3D and Superfamily (70). We predicted protein disordered regions using the PrDOS software (http://prdos.hgc.jp/about.html) (71).

#### Protein structure analysis

To construct 3D models of proteins, we searched their homologs in PDB database using BlastP, Delta-Blast and HHpred (70). Then, 3D structure models were built by homology modelling based on their homologous structures (PDB ID: 4xp4_A) using MODELLER (72). The quality of the models was assessed by Ramachandran plot analysis through PROCHECK (73). The images were generated with PyMOL software (http://pymol.org/) (74).

## RESULTS

### Genome sequencing of Schistocephalus solidus

#### Genome assembly and completeness

By combining short and long reads produced with Oxford Nanopore technology, we produced a genome assembly corresponding to 4 944 scaffolds. These scaffolds summed to over 500 Mb (80% of the assembled genome). To test the completeness of our assembly, we mapped all the available transcripts of *S. solidus* (33) on our *de novo* assembly. We found that more than 99.6% (24 676/24 765) of transcripts mapped onto the new assembly. The remaining 89 transcripts that were not found were blasted (blastn) against the assembly dataset. A total of 46 transcripts had a hit on 28 scaffolds but these hits were weak regarding the alignment length. This left 43 orphan transcripts that had no match in the assembly.

We also searched BUSCO groups to investigate the completeness of the genome assembly using a complementary approach. A total of 649 complete BUSCOS (66%) were found out of the 978 BUSCO groups in the metazoa database, which is close to the proportions obtained from the assembly of other cestode genomes (*e.g*. 72.6% in the genome of *Schistosoma japonicum* (79); 73.2% in the genome of *Schistosoma haematobium* (80)). The busco duplication rate was 0.8%. Overall, these results suggest that the coding genome was well-represented in the genome assembly. Finally, we identified a total of 56% of elements that appeared to be repeated in the new genome of *S. solidus* (Additional file 5), which is in accordance with previous studies that demonstrated that the genomes of most flatworms include large numbers of repetitive elements (81).

#### Number of putative genes

We first performed gene prediction using an *ab initio* approach, which gave a total of 21 780 ORFs. A second approach based on bam files from the transcript alignments of *S. solidus* (33) led to the identification of 30 103 ORFs. The two sets were merged and we found that 9 103 ORFs overlapped between the two sets, with a highly variable percent identity (ranging from 1% to 100%). Because of this variation in percent identity and in order to obtain exhaustive information, we decided to keep duplicate sequences for the rest of the analysis, which left a total of 51 883 ORF sequences identified. When comparing these predicted genes to the previously published transcriptome of *S. solidus*, we found that 77% of the transcripts were found in these predicted genes, and that 66% of the predicted genes were found in the transcriptome.

#### Identification of sequences specific to the new assembly

The 51 883 ORF sequences were aligned against a database of 43 058 protein sequences of *Schistocephalus solidus* obtained from NCBI. We identified 19 853 sequences that had no blast match and 16 287 sequences that had no significant match (E value < 1e-5). The total unmatched sequences were therefore 36 140. These unmatched sequences are potential protein coding genes not represented in public databases of *S. solidus* and were therefore used as one of the reference databases in the LC-MS/MS analysis.

#### Annotation of the new expanded *S. solidus* genome

The 51 883 ORF sequences were annotated using domain information (56) and orthology assignment (57). First, an important fraction of the predicted sequences (35 576 ORFs; 68.6%) did not have a match in any database and therefore functions could not be assigned to them. We refer to them as putative *S. solidus-*specific genes (Additional file 6). We found that 16 307 ORFs were successfully assigned a putative biological function (Additional file 6: 7 796 ORFs had a match on both databases, 4 591 ORFs had a match on the domain database only and 3 920 ORFs had a match on the orthology database only). Enrichment analyses on the new expanded *S. solidus* genome were conducted to identify potential class of genes that could be over-represented, and that could be either associated with *S. solidus* parasitic life style or with the phenotypic changes in the fish host. Overall, enrichment analyses showed that the functions were related to environment sensing: 2.5% of ORFs are *rd3* genes known to be expressed in photoreceptor cells (82); cell division, growth and development: 1.7% of ORFs encode for proteins involved in the movement of microtubules and 1.2% of ORFs are atrophin coding genes involved in development (83); as well as cell physiology: 1% of ORFs encode for PARP proteins involved in various cell physiological processes (84).

### Characterization of the proteome

Using mass spectrometry (LC-MS/MS), we detected 2 290 proteins in samples of whole *S. solidus* tissues, among which 1 467 proteins were detected in all worm samples (Additional file 7). Among these 2 290 proteins, 246 proteins were only detected using the new genome of *S. solidus* as a reference database during the LC-MS/MS analysis, with 113 proteins detected in all worms. The new genome sequence and annotation therefore provide a significant additional resource for *S. solidus* functional genomics. Most of the 2 290 proteins were functionally annotated, with the exception of the 246 proteins previously mentioned, for which functions were further investigated using a 3-step approach based on sequence, structure and phylogenetic analyses (see below). Using this approach, we found that the 2 290 proteins of *S. solidus* included 40 proteins encoded by 27 genes that did not have any sequence or domain similarities with other known species, and for which functions could not be described (*i.e*. 1.7 % of putative *S. solidus-*specific proteins in the proteome, compared to 68.6% of putative *S. solidus*-specific genes in the new genome) (Additional file 8).

Enrichment analyses in terms of biological process, cellular component and molecular function (GO terms) were performed for all the proteins detected in at least one of the worm samples, and also for the proteins detected in all worm samples. For proteins with several protein IDs, we kept only the first ID in order to prevent enrichment bias. We found that 14.9% of proteins detected in all worm samples were involved in protein metabolism processes, namely “translation” [GO:0006412] and “protein folding” [GO:0006457] (p<0.001). These are typically highly expressed proteins, which explains their over-representation here (81). Also, we found that 10.2% of proteins detected in all worm samples referred to mechanisms important in the functioning and the regulation of the cell cycle: “microtubule based process” [GO:0007017], with the microtubules having major roles in cell division and growth, and “tricarboxylic acid cycle” [GO:0006099], the latter referring to abundant metabolic enzymes that are involved in respiration by producing ATP (p<0.001) (Figure 2A). We also found that 14.4% of proteins detected in all worm samples were proteins with GTP binding [GO:0005525] and GTPase activity [GO:0003924] (p<0.001) (Figure 2B). Consistent with all the functions described above, we found that 23.2% of proteins detected in all worm samples were localized in the cytoplasm (p<0.001).

**Figure 2.**
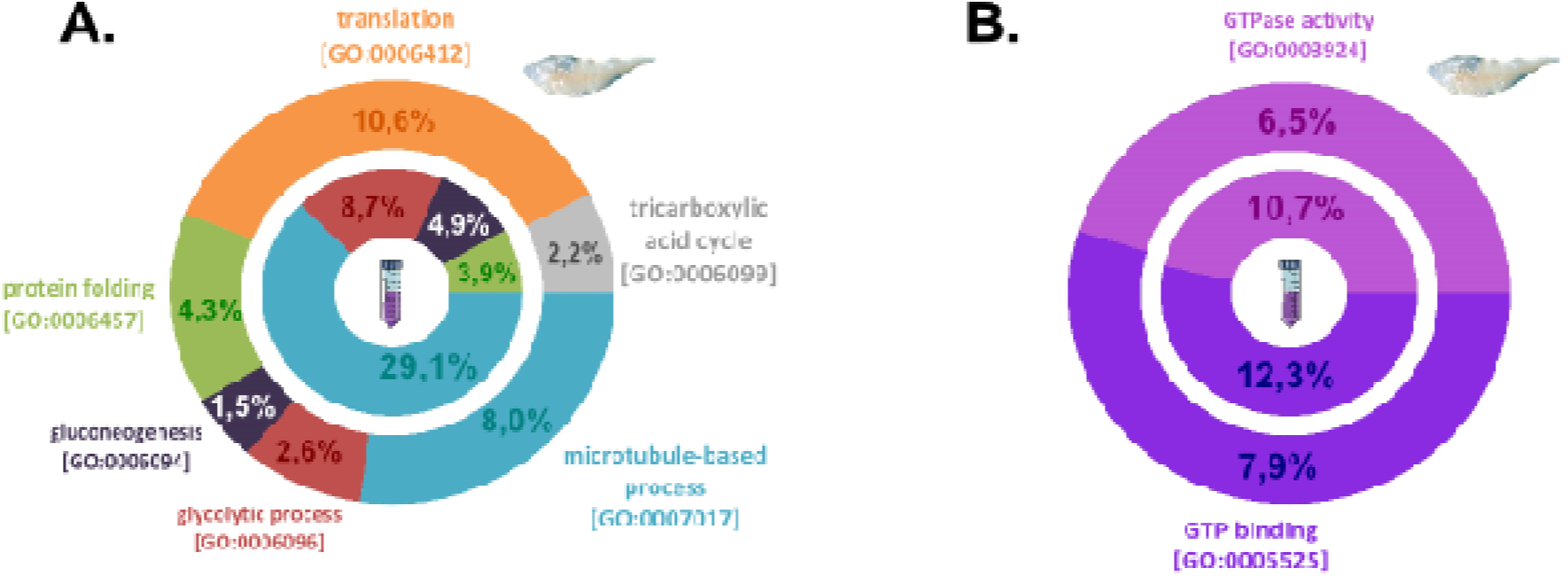
The proteome and secretome of *S. solidus* include proteins involved in protein metabolism/cell growth/energy intake. Results that were significant (p<0.001) for the enrichment analyses performed in terms of biological process (**A.**) and molecular function (**B.**). In each case, analysis was performed for the proteins detected in all worm (outer chart) and secretome (inner chart) samples.

The prediction of the functions of the 246 proteins that were detected using the new genome as a reference database was further investigated with sequence and structure analyses (Additional file 7). Their functions were in accordance with the previous enrichment results, as we found for example proteins involved in microtubules (protein ID: g20896.t1) or with ATPase activity (protein ID: g10622.t1). Additionally, we found that six proteins were peptidases or proteases with conserved active sites, suggesting that they are functional (Table 2), and that a protein contained an amidase domain. This protein “g7530.t1” is a Fatty acid amide hydrolase like (SsFAAH-like). FAAH enzymes degrade signalling lipids of the endocannabinoid class (94).

**Table 2.**
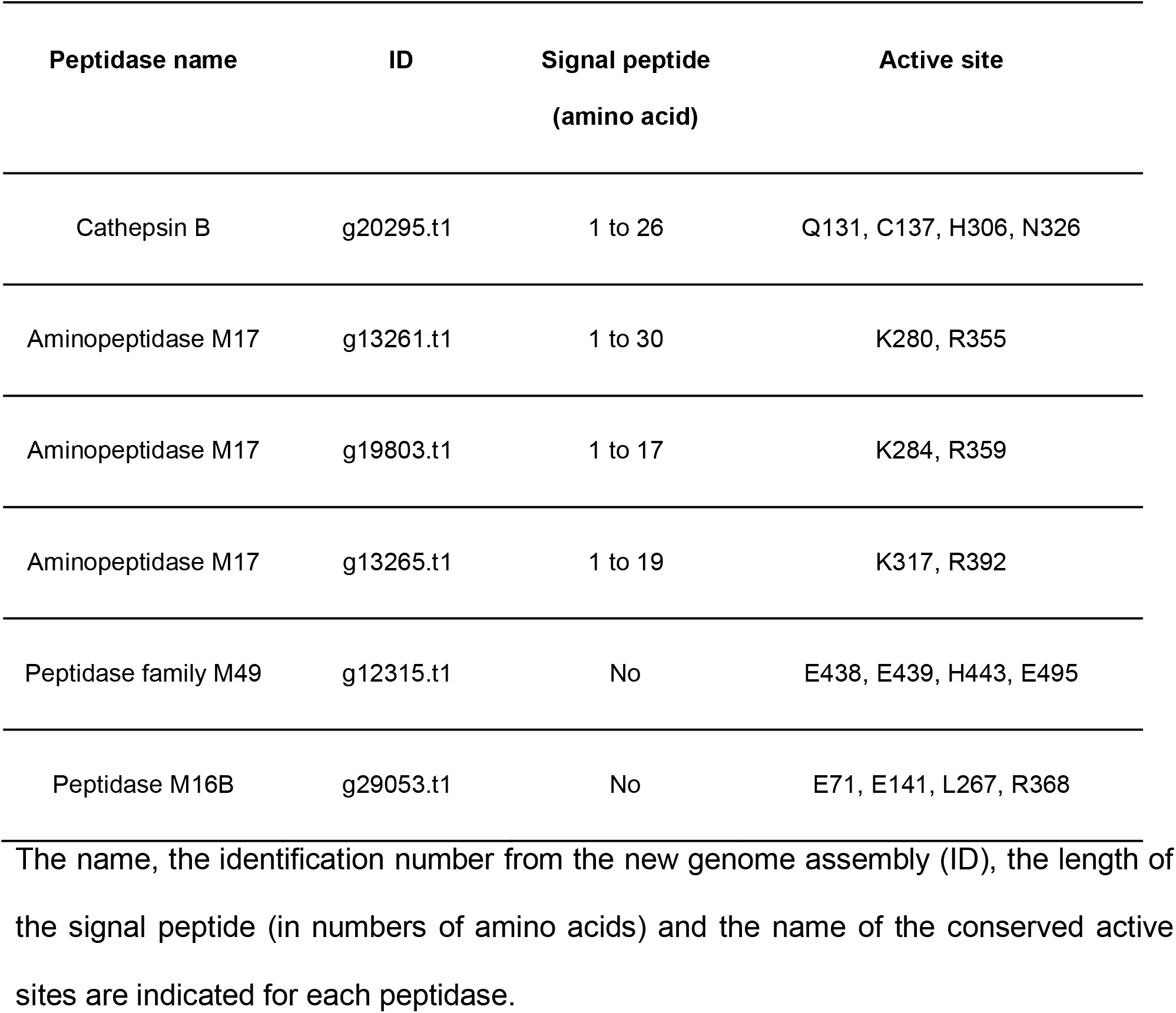
Peptidases detected in the proteome of S. solidus using its new expanded genome. The name, the identification number from the new genome assembly (ID), the length of the signal peptide (in numbers of amino acids) and the name of the conserved active sites are indicated for each peptidase.

| Peptidase name | ID | Signal peptide<br>(amino acid) | Active site |
| --- | --- | --- | --- |
| Cathepsin B | g20295.t1 | 1 to 26 | Q131, C137, H306, N326 |
| Aminopeptidase M17 | g13261.t1 | 1 to 30 | K280, R355 |
| Aminopeptidase M17 | g19803.t1 | 1 to 17 | K284, R359 |
| Aminopeptidase M17 | g13265.t1 | 1 to 19 | K317, R392 |
| Peptidase family M49 | g12315.t1 | No | E438, E439, H443, E495 |
| Peptidase M16B | g29053.t1 | No | E71, E141, L267, R368 |

Additionally, 141 proteins were assigned both to the worm and to the fish host during LC-MS/MS analysis (Additional file 9). As it was not possible to directly determine if the 141 proteins were produced by the worm or the fish, we searched if these proteins could be core proteins, *i.e.* proteins conserved in all eukaryotes and whose functions are well characterized (85). Using Blastp and Pfam tools, we found that the 141 proteins had sequence similarities with 248 previously reported eukaryotic core proteins (85) (mean Expect-value 2.45e-11), and that they were composed of conserved domains. We therefore concluded that these proteins were expressed both in the worm and in the fish because of their fundamental functions in the cell but that their peptide similarities did not allow to determine their origin. Enrichment analysis performed on these 141 proteins confirmed that they were involved in cell physiology and in energy production as found for the whole proteome.

### Characterization of the secretome

A total of 1 568 proteins were detected in the secretome samples (Additional file 7). The numbers ranged between 781 and 1 183 proteins depending on the sample, with 459 proteins detected in all secretome samples. As expected, the secretome of *S. solidus* was mostly a subset of the worm proteome, both in terms of protein number and functions, with a few exceptions described below. The whole protein content of the secretome of a given worm represented up to 59% of the whole protein content of the proteome of that same worm. In total, 1 538 unique proteins were shared between at least one secretome sample and one proteome sample, among which 385 proteins were detected in all secretomes and proteomes studied, and 74 proteins were detected in all secretomes and in at least one proteome. All the proteins that were shared between secretome and proteome samples had a functional annotation, except for 36 proteins that were previously detected in the proteome as *S. solidus*-specific and that were also secreted (2.3% of putative *S. solidus*-specific proteins in the secretome, compared to 68.6% of putative *S. solidus*-specific genes in the new genome and 1.7% of putative *S. solidus*-specific proteins in the proteome) (Additional file 8).

We found that the secretome was composed of proteins enriched for biological processes, cellular components and molecular functions that were similar to the proteome. For the proteins detected in all secretomes, almost half of the proteins participated in the regulation of cell division and energy production. These proteins were enriched in functions such as “microtubule based process” [GO:0007017] involved in the regulation of cell division, “glycolytic process” [GO:0006096] and “gluconeogenesis” [GO:0006094], these two processes being important source of energy production (p<0.001 hypergeometric test corrected with Bonferroni method) (Figure 2A). Proteins were also enriched in domains with GTP binding [GO:0005525] and GTPase activity [GO:0003924] (Figure 2B), as previously reported in the proteome (p<0.001). Furthermore, 37.9% of the proteins detected in all the secretome samples were predicted to be localized in the cytoplasm (p<0.001), similar to proteins in the proteome. Notably, we found that a significant number of proteins (6.1% of the proteins detected in all secretomes p<0.001) were reported to be specifically localized into the extracellular space, an enrichment that was specific to the secretome.

### Proteins unique to the secretome

We found eight proteins that were detected in all secretome samples, but in none of the proteomes (Table 3). Among them, three proteins had fibronectin type-III domains. The first (protein ID: A0A0X3PH69) was a “Neogenin”, which is a protein involved in neural development with two fibronectin type-III domains. The second (protein ID: A0A0X3Q1B7; A0A0X3PKA1; A0A0X3Q8R6) was a Receptor-type tyrosine-protein phosphatase eta that included three fibronectin type-III domains. The third (protein ID: A0A0V0JBL5) included two fibronectin type-III domains, but no additional information on the function of this protein was available. Furthermore, we detected an uncharacterized protein with a predicted molecular function corresponding to “Neurotransmitter: sodium symporter activity (NSS)”, which is involved in transmembrane transport (protein ID: A0A183S8K9; A0A0X3PDV9; A0A0X3PNC8; A0A0V0J682; A0A183T7R5), and a protein potentially acting at the cell membrane as a Phospholipid scramblase (PLSCR) (protein ID: A0A0X3P711; A0A183SGM7). Lastly, three other uncharacterized proteins did not have information on function using Uniprot database, Blast tools and protein domain identification. It seems that these three proteins are specific to *S. solidus*, with the last protein (protein ID: A0A0X3Q756) having a secretory signal peptide.

**Table 3.** Proteins that are excreted/secreted by *S. solidus*, detected in five.

| Uniprot ID | Information from Uniprot and Blast tools | Information from sequence, structure and phylogeny |
| --- | --- | --- |
| A0A0X3PH69 | Neogenin | Fibronectin type-III domains |
| A0A0X3Q1B7;A0A0X3PKA1;A0A0X3Q8R6 | Receptor-type tyrosine-protein phosphatase eta | Fibronectin type-III domains |
| A0A0V0JBL5 | Unknown | Fibronectin type-II domains |
| A0A183S8K9;A0A0X3PDV9;A0A0X3PNC8;A0A0V0J682;A0A183T7R5 | Neurotransmitter: sodium symporter (NSS) | Unknown |
| A0A0X3P711;A0A183SGM7 | Phospholipid scramblase | Unknown |
| A0A0V0J8W2 | Unknown | Gene specific to *S. solidus* (orphan) |
| A0A0X3P740 | Unknown | Gene specific to *S. solidus* (orphan) |
| A0A0X3Q756 | Unknown | Gene specific to *S. solidus* (orphan)<br>Signal peptide (secreted) |

Protein IDs were taken from Uniprot. When several protein IDs were assigned to one protein, these protein IDs corresponded to isoforms with identical functions. For each protein, functional annotation was first retrieved from searches with Uniprot and Blast tools. Complementary analyses based on sequence, structure and phylogeny were used to obtain information.

Twenty-two proteins were detected in at least one of the five secretome samples, but in none of the proteomes (Table 4). Three proteins were composed of fibronectin type-III domains. The first (protein ID: A0A0X3NX35) detected in four out of five secretomes was described as a Receptor-type tyrosine-protein phosphatase H. The second (protein ID: A0A183TLI3; A0A0X3PWG6; A0A0X3PW04; A0A0V0J316; A0A0X3PN23; A0A0X3PJY8), which was detected in three secretomes, was either a Tenascin (an extracellular matrix glycoprotein) or a Receptor-type tyrosine-protein phosphatase F. The last protein of this type (protein ID: A0A0V0JA38), also detected in three secretomes, was, according to the blast result, a collagen-like protein, which is an important component of cuticle.

**Table 4.**
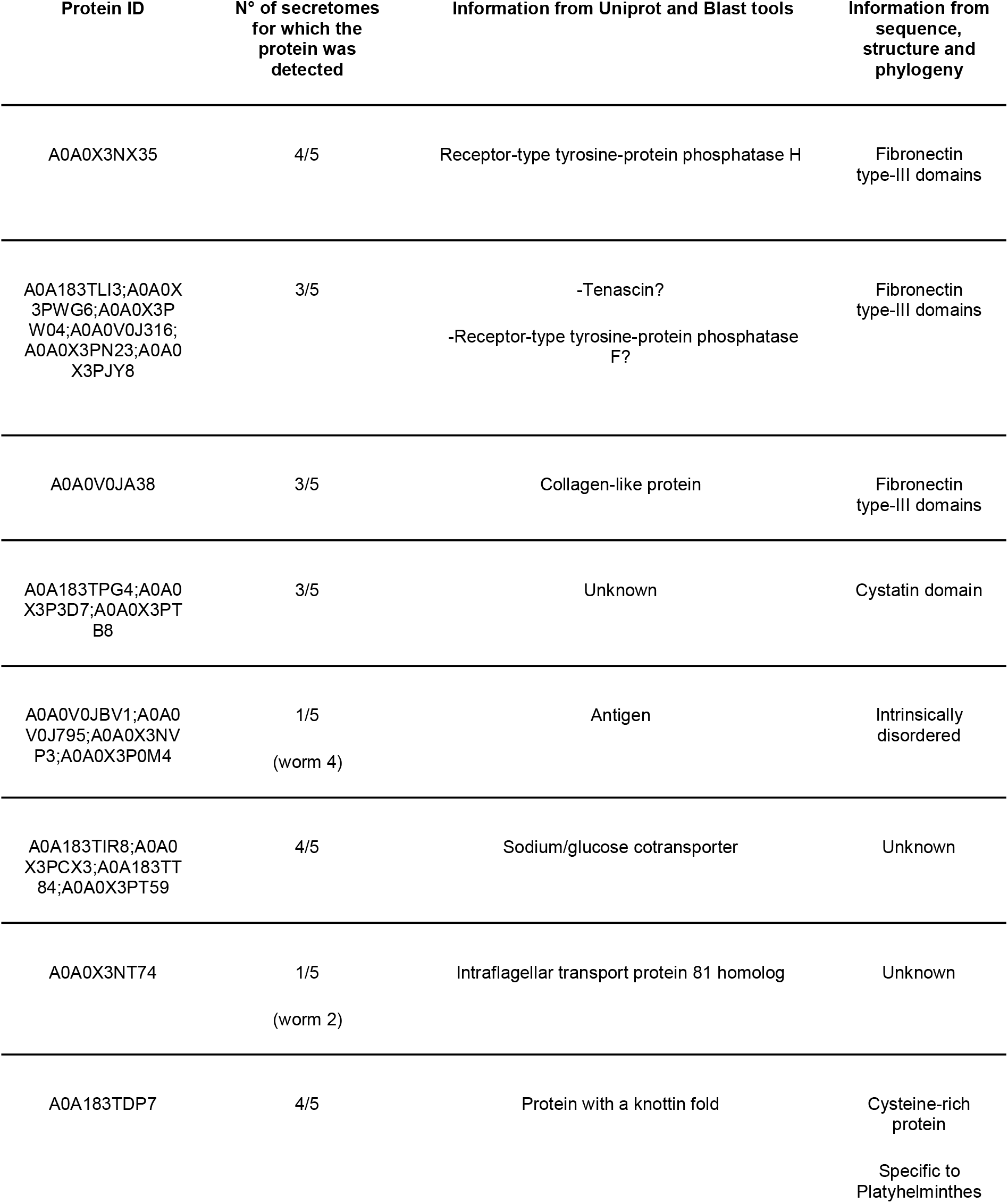
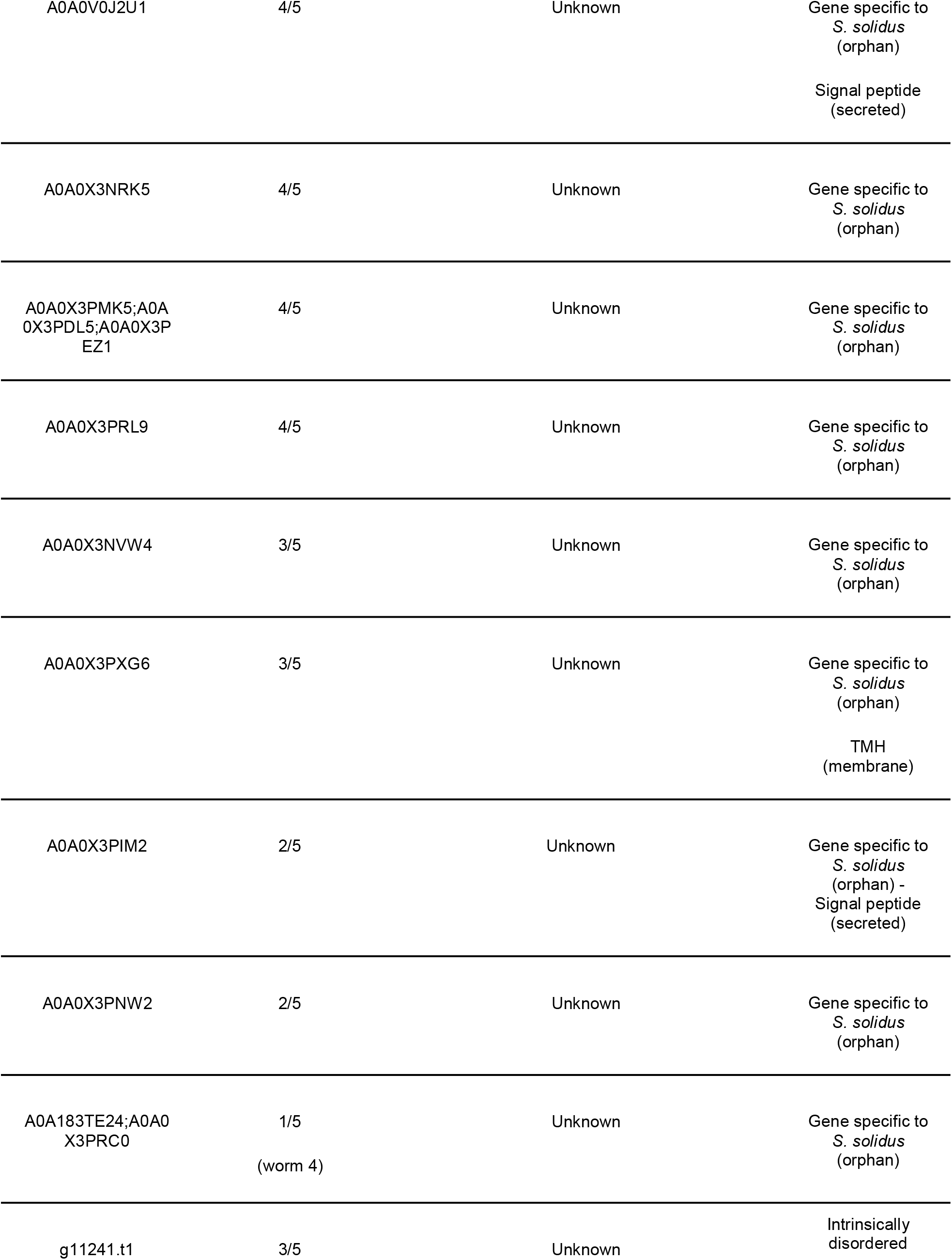
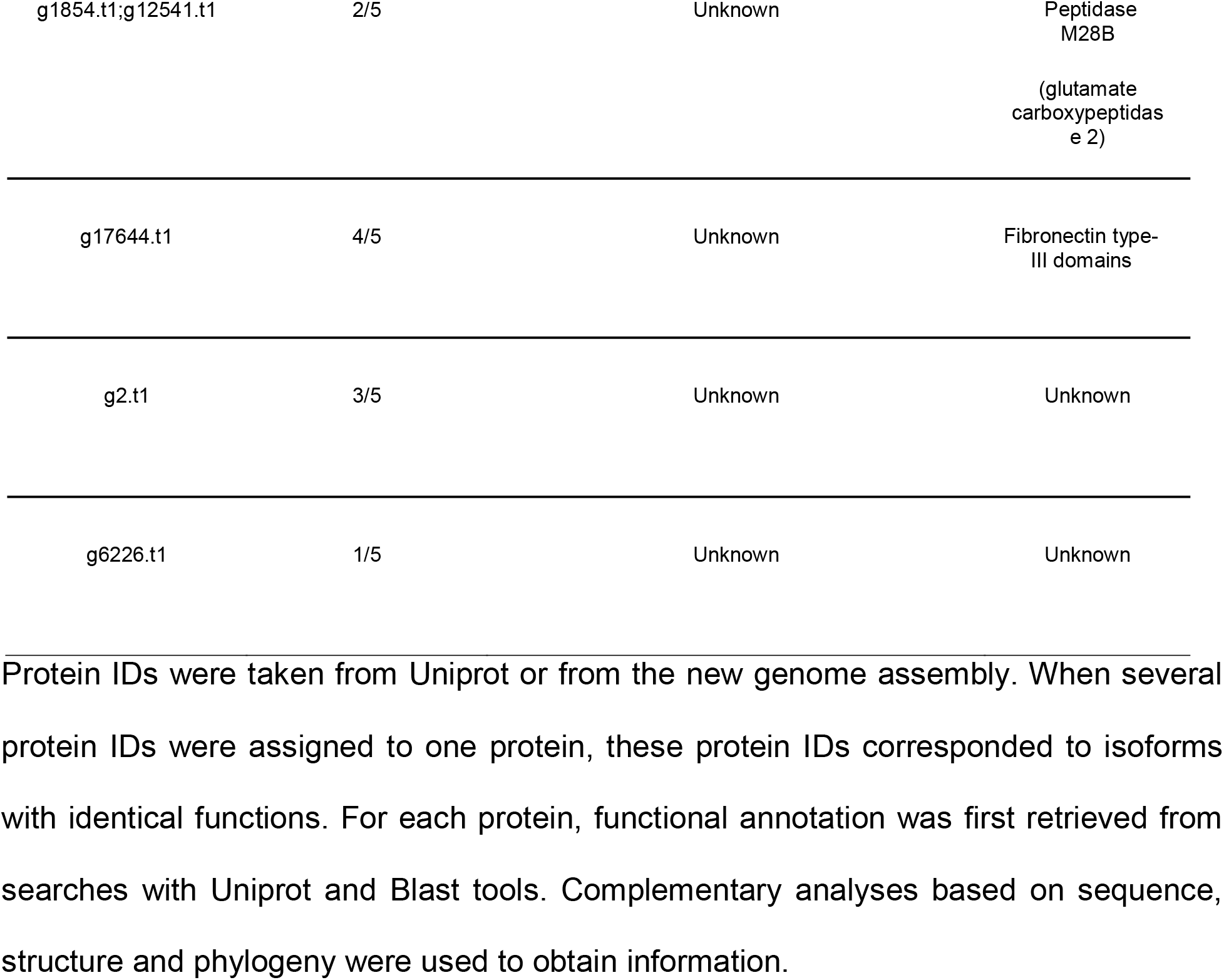
Proteins that are excreted/secreted by *S. solidus*, detected in at least one secretome and not in proteomes.

Two proteins appeared to be involved in immunity processes. The first (protein ID: A0A183TPG4;A0A0X3P3D7;A0A0X3PTB8), which was detected in three secretomes, was uncharacterized and had a cystatin domain which may be involved in immuno-modulatory functions. The second (protein ID: A0A0V0JBV1; A0A0V0J795; A0A0X3NVP3; A0A0X3P0M4), which was detected in the secretome of the largest worm only (worm 4), was according to the blast results an antigen similar to the diagnostic antigen gp50 commonly used to detect parasitic diseases.

Two proteins were associated with transport functions. The first (protein ID: A0A183TIR8; A0A0X3PCX3; A0A183TT84; A0A0X3PT59) was detected in four out of five secretomes and was a sodium/glucose cotransporter involved in glucose homeostasis. The second (protein ID: A0A0X3NT74), which was only detected in the second largest worm (worm 2), was an intraflagellar transport protein required for ciliogenesis.

One protein (protein ID: A0A183TDP7), detected in four secretomes, had a knottin fold similar to a domain found in the *Schistosoma* parasitic trematodes, but for which the function is not characterized. It is a cysteine-rich protein and interestingly, blastP search against UniProt database revealed that it is only found in platyhelminthes and amino acid multiple sequence alignment showed that it presents a new motif, C-x(6)-C-x(7)-CC-x(4)-C-x(9)-C-x(2)-C-x(6)-C-x(5)-CC-x(3)-C-x(4)-C. This type of protein is resistant to proteases.

Nine proteins had functions which could not be clearly determined by blast in Uniprot and Non-redundant protein sequences (GenBank) databases. Five other proteins were only detected using the new genome of *S. solidus* as a database reference. We further investigated the functions of these last 14 proteins with sequence, structure and phylogenetic analysis: nine proteins were specific to *S. solidus*. Among them, two proteins (protein ID: A0A0V0J2U1 - protein ID: A0A0X3PIM2) had secretion signal peptides and one protein (protein ID: A0A0X3PXG6) had a TMH (Trans-Membrane Helix) domain found in transmembrane proteins. Furthermore, one protein was identified as a peptidase M28B (glutamate carboxypeptidase 2) (protein ID: g1854.t1; g12541.t1) and one protein had fibronectin type-III domains (protein ID: g17644.t1).

## DISCUSSION

Understanding the molecular mechanisms behind behavioural changes in a vertebrate host infected by a non-cerebral parasite is a fascinating challenge. If these changes are the result of parasite manipulation of its host, one possible mechanism hinges on molecules secreted by the parasite (*i.e.* manipulation factors) that impact the host physiological, immunological and central nervous systems, and ultimately the behaviour of the host. Here, we describe for the first time the protein component of the secretome of a parasite commonly referred as manipulative, using mass spectrometry (LC-MS/MS), with the objective to identify potential manipulation factors that could explain host behavioural changes. As expected, we found that the proteins that are excreted/secreted by *S. solidus* are mostly a subset of the proteins being expressed in the whole worm, and that both included proteases. We also found that 30 secretome proteins were not detected in the proteome and were involved in neural, immune, and cell communication functions, therefore having the potential to interfere with the host physiological systems and behaviour. Finally, we highlighted that the secretome of *S. solidus* included *S. solidus*-specific proteins that could play important roles in the tight interaction of the parasite with its fish host. All together, these proteins represent promising candidates to explain physiological and behavioural changes in the stickleback host.

### The secretome of *S. solidus* is mostly a subset of its proteome with similar functions

The global protein composition of the proteomes and the secretomes of *S. solidus* highlights functions that are crucial for its parasitic lifestyle and for its interactions with the external environment. As it is generally reported (39), we found that the proteins that are excreted/secreted by *S. solidus* (*i.e.* the secretome) are mostly a subset of the proteins being expressed in the whole worm (*i.e.* the proteome), both in terms of protein number and function. We found that both the proteome and the secretome of *S. solidus* at the infective stage (*i.e.* when behavioural changes appear in the fish host) were enriched in proteins involved in cell division, which was also found in the functional analysis of the genome we sequenced, as well as energy production. Concerning cell growth, our results may seem counter intuitive because a previous transcriptome analysis revealed that the non-infective stage of *S. solidus* shows higher expression levels of genes involved in growth and cell regulatory functions compared to the infective stage (33). However, we did not analyse the proteome and the secretome of worms at the non-infective stage, such that we can only quantify that these processes are enriched at the infective stage but not their relative importance compared to other life stages of the parasite. We therefore only speculate that both the non-infective and the infective stages of *S. solidus* could rely on biological processes involving cell growth, but production levels would be much higher at the non-infective stage, considering the fact that growth occurs predominantly in the first 12 weeks after installation in the fish host (86). Concerning energy production, the results are also surprising, as the worm was empirically described to use its glycogen reserve (its primary source of energy), mainly when it reaches its final avian host (87). However, transcriptomic analyses demonstrated that glycogen metabolism and energy production in *S. solidus* are complex processes (33). Specifically, 6 steps of the glycolysis cycle are highly expressed at the infective stage (33). Furthermore, it is important to keep in mind that, for tissue and secretome samples collection, worms were placed in a solution of Phosphate Buffered Saline (PBS, pH 7.4) in the dark for 2 hours. While this method is commonly used to collect parasitic secretomes (59), we cannot dismiss the possibility that the environmental change (from the body cavity of the stickleback to the PBS tube) encountered by *S. solidus* at this step could have influenced the levels of proteins detected in the proteomes and secretomes. One can speculate that this environmental stress could have triggered cell growth and energy intake in the worm (for example to counteract other metabolic effects), explaining the discrepancies between the proteomics data we obtained and the previous transcriptomic work (33). To confirm our results, future work could replace the PBS solution by a medium that would better mimic the fish abdominal cavity, for example by a medium already used *in vitro* for breeding (21). The global analysis of the proteome and of the secretome of *S. solidus* therefore confirms the importance of energy use for the worm, even before it reaches the bird.

We detected both in the proteome and the secretome of *S. solidus* proteins with GTP binding and GTPase activity. GTP binding proteins also called G-proteins are known to regulate a variety of biological processes such as mediating signals by hormones and light, gene expression, cytoskeletal and microtubule organization, or vesicle trafficking (88–91). In parasites, GTPases have been demonstrated to have important roles in the secretion of virulence factors (90, 91). For instance, in *Toxoplasma gondii*, which is a parasite known to induce behavioural changes in rodents (92), Rab GTPases regulate the secretion of proteins essential to invade host cells, and the modification of their expression results in aberrant transport of proteins (93).

The use of the new genome of *S. solidus* as a reference database during mass spectrometry analysis allowed us to detect 6 peptidases or proteases in the proteome of *S. solidus*, and one peptidase (peptidase M28B glutamate carboxypeptidase 2) detected only in the secretome fraction, with proteases and peptidases being typically described as virulence factors for many parasites (Table 1). In *S. solidus*, proteases and peptidases may have important roles in weakening the fish immune response at the infective stage when the worm is ready to pass into its avian host to reproduce (21). Injecting these proteases and peptidases alone or in combination, into isolated head kidney samples from non-infected fish as in (21), would be necessary to confirm their potential role in disrupting the host immunity. Furthermore, this shows that the new genome presented here is a valuable tool to identify proteins that are critical for the parasitic lifestyle. We also found an SsFAAH-like enzyme that could potentially degrade signalling lipids of the endocannabinoid class. Endocannabinoids have previously been reported as an important player in host-parasite interactions, by promoting the activation of the immune response in the host (94–95). This enzyme was not detected in the secretome of *S. solidus*.

### The secretome of *S. solidus* also contains proteins not detected in the proteome

#### Neuronal and immune functions

Transcriptomic analysis demonstrated that when *S. solidus* reaches the infective stage, genes involved in neural pathways and sensory perception are expressed by the worm at higher levels (33). Thus, we expected the secretome at the infective stage of *S. solidus* to be enriched with proteins involved in such neural functions (as described in other parasitic systems, reviewed in Table 1). Furthermore, behavioural changes in the stickleback infected by *S. solidus* generally appear in concomitance with the activation of the immune response of the fish (21). Therefore, we expected the secretome of *S. solidus* at the infective stage to include proteins involved in immunity (as reviewed in Table 1). We found that three proteins were playing potential roles in neural and immune functions. The first protein, which was detected in four secretomes but in none of the proteomes, had a knottin fold called UPF0506 composed of cysteines and generally found in *Schistosoma* parasites. In *Schistosoma*, the function of the proteins with such knottin fold is not defined. However, peptides with knottin domains (*i.e.* knottins) were described in venoms from various animals. For venomous animals, knottins are neurotoxins having high specificity towards receptors in the nervous system of their prey or aggressor (96). Furthermore, it is a cysteine-rich protein, and in parasites cysteine-rich proteins play a role in invasion (97) and modulation (98) of the immune system. Therefore, this protein could be a promising manipulation factor if it could act as a neurotoxin in the brain of infected sticklebacks. The second protein, which was detected in three secretomes but in none of the proteomes, had a cystatin domain which may be involved in immuno-modulatory functions. It was demonstrated in parasitic nematodes that cystastins are important secreted molecules that help parasites to evade immunity of the host (99). Nematode cystatins inhibit host proteases involved in antigen processing and presentation, leading to a reduction of the host immune response (100). This secreted protein could thereby explain in part why the immune response is only activated late in infected sticklebacks, which needs to be further studied. The third protein appeared to be an antigen similar to the diagnostic antigen gp50. This diagnostic antigen is used to detect parasitic infection, for example by *Taenia solium* (101). However, the antigen was detected only in the secretome from the biggest worm of our study and not in its proteome. Sampling over a longer time frame could alleviate this type of discrepancies between samples. One protein, which was detected in all secretomes but in none of the proteomes, had a “Neurotransmitter: sodium symporter activity (NSS)” annotation. In humans, it is a membrane protein involved in the termination of synaptic transmission and the recycling of neurotransmitters at the membrane (102). How this membrane protein could be secreted and act as a manipulation factor in the secretome is unclear. Therefore, we are cautious in interpreting the role of this protein as a manipulation factor as it could be solely a waste product from the membrane of the parasite.

#### Cell communication

*S. solidus* is located in the abdominal cavity of the threespine stickleback (16). As the worm is not in direct contact with its host brain, we expected to detect proteins involved in cell communication or cell-cell signalling. During annotation of the new genome, 2.5% of genes were related to environment sensing functions. Furthermore, we found that 7 proteins detected only in the secretome fraction were characterized by fibronectin type-III (FNIII) domains. FNIII domains are widely found in animal proteins and are involved in cell-cell interactions (103). The first protein with fibronectin type-III domains found in all secretomes but in none of the proteomes was identified as a Neogenin. In addition to playing roles in cell-cell adhesion, neogenins are involved in neural development in humans (104). Three additional proteins that we identified with fibronectin type-III domains were described as Receptor-type tyrosine-protein phosphatases (type eta, detected in all secretomes but in none of the proteomes, type H, detected in 4 secretomes but in none of the proteomes, type F, detected in 3 secretomes but in none of the proteomes). Receptor-type tyrosine-protein phosphatases are involved in cell-cell communication (105). It was previously shown that a baculovirus secretes a protein tyrosine phosphatase, which acts on the neural system of its host the silkworm *Bombyx mori* and enhances its locomotory activity, so that it ultimately increases the virus dispersal (106) (Table 1). The set of phosphatases identified here are located in the membrane, such that they are predicted to have different specific biological functions that the protein tyrosine phosphatase found in the baculovirus. However, their abundant representation in the secretome is intriguing and the importance of the process of phosphorylation represents a future avenue of study in the context of behavioural changes in the fish host, such as increased exploration (23). For the next two proteins identified with fibronectin type-III domains, the first protein was found in all secretomes but in none of the proteomes and had no clear function (protein ID: A0A0V0JBL5), as for the second protein that was detected in 4 secretomes but in none of the proteomes (protein ID: g17644.t1). The last protein appeared to be a collagen-like protein found in 3 secretomes. In *Caenorhabditis elegans*, the cuticle is an extracellular matrix made of like proteins (107). Thus, the role of this protein in cell communication is uncertain and it may rather be a waste product excreted passively from the cuticle of *S. solidus*. In summary, fibronectin type-III proteins secreted by *S. solidus* appear to be good candidates for manipulation factors because of their roles in cell-cell signalling, but also in potential neural functions.

Additionally, we expected to detect proteins with transport functions in the secretome of *S. solidus* to mediate the communication between the worm localized in the host abdominal cavity and the host brain. We detected three proteins involved in transport or related functions at the membrane, including a protein that was detected in all secretomes but in none of the proteomes, a Phospholipid scramblase. It is a transmembrane protein that is known in human to bind to the 5’-promoter region of the *inositol 1,4,5-triphosphate receptor type 1* gene (*IP3R1*) so that it enhances expression of the receptor (108). Since these proteins detected here could be simply waste products coming from the membranes of the parasite, we thus have to be careful when speculating about the roles of these proteins, and functional analyses will be required.

### The secretome includes *S. solidus*-specific proteins that could play important roles in parasite-host interactions

During annotation of the new genome, we found that 68.6% of the predicted genes did not have any sequences or domain similarities with other known species that would allow accurate annotation, therefore representing putative *S. solidus*-specific genes. Part of these genes are effectively translated, as 1.7% of the proteome and 2.3% of the secretome included putative proteins specific to the worm. Specifically, we detected a total of 12 proteins only in the secretome fraction for which functions could not be clearly determined, but that were specific to *S. solidus*. Three proteins were detected to be secreted signal peptides (protein ID: A0A0X3Q756 - protein ID: A0A0V0J2U1 - protein ID: A0A0X3PIM2). We found that the proteins that are specifically enriched in the secretome are more likely to have secreted signal peptides (10% of the proteins detected only in the secretome fraction had secreted signal peptides, while 0.17% of the proteins detected in the proteome had such signals). One protein had a TMH (Trans Membrane Helix) domain found in transmembrane proteins (protein ID: A0A0X3PXG6). Proteins with TMH domains are used to diagnose parasitic infection as they are highly specific to the parasite of interest (110, 111). Interestingly, analyses conducted with the transcriptome of *S. solidus* previously suggested that 19% of all the protein coding genes could be *S. solidus*-specific (32). In this study, producing a novel genome sequence and assembly of *S. solidus* led us to increase this estimation by three times. *S. solidus*-specific secreted proteins represent promising candidates to explain physiological and behavioural changes in the stickleback host. The threespine stickleback is an obligatory host of S*. solidus* (112). Because of the potential co-evolution between the worm and the fish (16), *S. solidus* may have developed highly specific molecular mechanisms targeting the threespine stickleback physiological machinery to insure it can establish and grow in this fish only, until it is ready to pass into its final avian host. The *S. solidus*-specific proteins found here are therefore likely to be part of this unique set of molecules crucial for the survival of the worm and we can assign them this ecological annotation (113). It will be very interesting in the future to produce recombinant *S. solidus*-specific proteins and test their effects on the phenotype of the threespine stickleback by functional analysis. These proteins could also serve to obtain structural information by X-ray crystallography or NMR spectroscopy, which could help us to discover their function. Finally, studies of how these proteins evolve between *Schistocephalus* populations and between species of *Schistocephalus* infecting different fish species (123) will shed additional light on the specificity and potential function of these proteins.

## CONCLUSIONS

The secretome of *S. solidus* appears to be an important component of the molecular interaction between the parasite and its threespine stickleback host. In accordance with our predictions, we detected in the secretome of *S. solidus* at the infective stage (*i.e.* when behavioural changes appear in the stickleback) proteases, proteins involved in neural and immune functions, as well as proteins involved in cell communication. We also highlighted in the secretome the presence of *S. solidus*-specific proteins. In the future, comparative studies could be conducted to validate that the proteins detected in the secretome of *S. solidus* act as manipulation factors in the behaviour of its fish host: at the organism level, the analysis of the secretome of the worm at the non-infective stage will confirm if the putative manipulative proteins reported in this study are effectively detected solely at the infective stage (or at highest levels) when the behavioural changes appear in the host. It would also be of special interest to study the secretome of *S. solidus* at the stage when it is able to reproduce, *i.e*. in the bird host. At the species level, comparing the secretome of *Schistocephalus solidus* with the secretome of a close related species *Schistocephalus pungitii* will be of great interest as *S. pungitii* does not have known effects on the behaviour of the nine-spine stickleback, its specific intermediate host species (114). Finally, functional studies testing effects of presence in non-infected sticklebacks of interesting proteins identified in the secretome, such as Receptor-type tyrosine-protein phosphatases or Phospholipid scramblase, would allow to better understand the contribution of these proteins in the behavioural changes. To conclude, we hope that the genomic and proteomic resources we provide will help other researchers to investigate general questions on host-parasite interactions in nature.

## DECLARATIONS

### Ethics approval and consent to participate

This study was approved by the Comité de Protection des Animaux de l’Université Laval (CPAUL 2014069-3).

### Consent for publication

Not applicable

### Availability of data and materials

The datasets supporting the conclusions of this article are included within the article and its additional files. Furthermore, the dataset of the new genome assembly is available from the NCBI BioProject with accession PRJNA576252. The mass spectrometry proteomics data have been deposited to the ProteomeXchange Consortium via the PRIDE partner repository with the dataset identifier PXD016844.

### Authors’ contributions

C.S.B. designed the study with input from N.A.-H., C.R.L., L.J.F. and K-M.M. C.S.B. performed the experiments to prepare the worms and their secretomes for mass spectrometry. K-M.M. performed the experiments with the mass spectrophotometer and the MaxQuant software. H. Martin prepared the worm samples for genome sequencing and performed the MinION sequencing. J.L. analyzed the genome and prepared the files used for searches with the MaxQuant software. J.L. and H. Maaroufi assembled and annotated the new genome and proteins. C.S.B. analyzed the data with input from N.A.-H., C.R.L. and L.J.F. C.S.B. and N.A-H. wrote the manuscript with input from all authors. All authors read and approved the final manuscript.

## Acknowledgments

We thank Karina Nielsen, Alexandra Sébastien and Tom Clark from the Foster lab for their advice during the preparation of the samples for mass spectrometry analysis. The authors thank the personnel of the Laboratoire Aquatique de Recherche en Sciences Environnementales et Médicales at Université Laval for their help with fish rearing. We thank Eric Normandeau from Ressources Aquatiques Québec (RAQ) for his help to identify *S. solidus* specific proteins.

## ADDITIONAL INFORMATION

**Additional file 1.** Visualization on SDS-PAGE gels of the protein content of the proteome and of the secretome for 5 *S. solidus* worms. A. Proteome (*i.e.* tissues) extracts from 5 distinct worms. Note that for worm 1 and worm 2, the sample was loaded on gel either in duplicate or triplicate but only one lane was used for LC-MS/MS. **B.** Secretome extracts from 5 distinct worms. On each gel, a BenchMark protein ladder (Invitrogen) and a negative control (SDS-PAGE loading buffer and water) were also loaded.

**Additional file 2**. Methods information for LC/MS analysis.

**Additional file 3**. Template custom script for analysis of the proteome and secretome LC-MS/MS data using Python.

**Additional file 4.** List of the proteins that were solely attributed to the threespine stickleback host, and that were consequently discarded from analysis. All the protein IDs that were specific to the threespine stickleback *Gasterosteus aculeatus* are reported. For a proteome or a secretome sample, a value above 0 indicates protein detection. The Gene Ontology (GO) annotation (biological process, cellular component and molecular function) of each protein ID is reported.

**Additional file 5.** List of the repeated elements that were identified in the new genome of *S. solidus*. For each class of repeated elements are indicated: the number of elements, their total length (pb) as well as their proportion (%) in the new genome.

**Additional file 6.** Annotation results of 16 307 ORFs predicted from the new expanded *Schistocephalus solidus* genome. Functional annotation was performed using PFAM (domain database) and eggNOG (orthology database). For each ORF match in each database: the accession number, the E-value and the functional description of the match are indicated. The other ORFs (68.6%) predicted from the new genome did not match with these two databases, and are considered *S. solidus*-specific.

**Additional file 7.** List of the proteins detected by LC-MS/MS in the proteome and the secretome of *S. solidus.* All the protein IDs that were detected in each of the proteome samples and/or secretome samples are reported. For a sample, a value above 0 indicates protein detection. The Gene Ontology (GO) annotation (biological process, cellular component and molecular function) of each protein ID is reported.

**Additional file 8.** List of the *S. solidus*-specific proteins detected in the proteome and/or in the secretome of *S. solidus*. For each protein, the ID is reported from Uniprot. If the protein is “not secreted”, then it was only detected in the proteome and not in the secretome. The Gene Ontology (GO) annotation (biological process, cellular component and molecular function) of each protein ID is reported.

**Additional file 9.** List of the 141 proteins that were assigned both to *S. solidus* and to the fish host during LC-MS/MS analysis. Each protein is assigned several protein IDs from the worm and the fish. The Gene Ontology (GO) annotation (biological process, cellular component and molecular function) of each protein ID is reported.

