## Supplementary figures and images for "The parasite *Schistocephalus solidus* secretes proteins with putative host manipulation functions"

### Additional file 1

#### A. Proteome of *S. solidus*

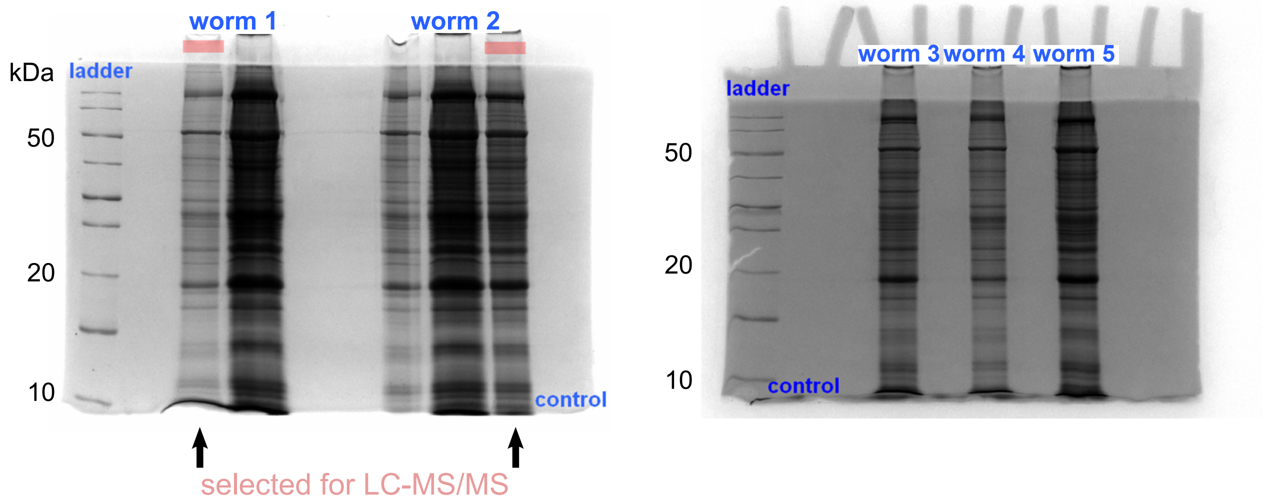


#### B. Secretome of *S. solidus*

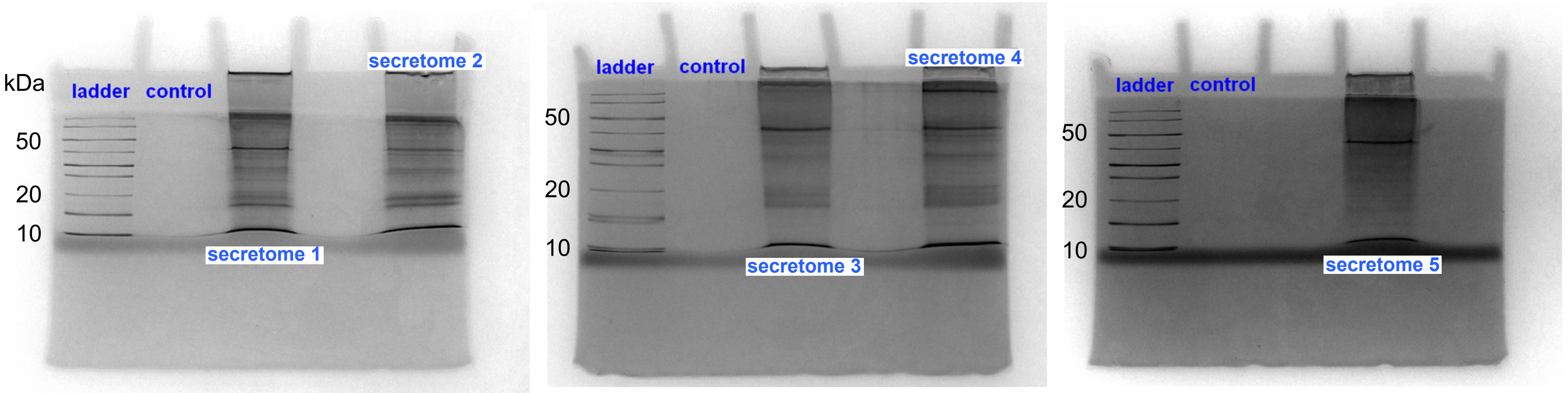
