## Additional file 2 for "The parasite *Schistocephalus solidus* secretes proteins with putative host manipulation functions"

LC-MS/MS analysis

For each sample, peptides obtained at the end of the STAGE-TIP protocol were dissolved in 0.1% formic acid with 2% acetonitrile. Peptides were analyzed by a quadrupole–time of flight mass spectrometer (Impact II; Bruker Daltonics) coupled to an Easy nano LC 1000 HPLC (ThermoFisher Scientific) using a 40–50 cm analytical column long. We used a 75-μm inner diameter fused silica with an integrated spray tip pulled with P-2000 laser puller (Sutter Instruments), packed with 1.9 μm diameter Reprosil-Pur C-18-AQ beads (Maisch, www.Dr-Maisch.com), and operated at 50°C with in-house built column heater. Buffer A consisted of 0.1% aqueous formic acid, and buffer B consisted of 0.1% formic acid and 80% (vol/vol) acetonitrile in water. A standard 90-min peptide separation was done, and the column was washed with 100% buffer B before re-equilibration with buffer A. The Impact II was set to acquire in a data-dependent auto-MS/MS mode with inactive focus fragmenting the 20 most abundant ions (one at the time at a 18-Hz rate) after each full-range scan from m/z 200 to m/z 2,000 at 5 Hz rate. The isolation window for MS/MS was 2–3 depending on the parent ion mass to charge ratio, and the collision energy ranged from 23 to 65 eV depending on ion mass and charge. Parent ions were then excluded from MS/MS for the next 0.4 min and reconsidered if their intensity increased more than five times. Singly charged ions were excluded from fragmentation (64).
